# The effect of the number motor units and their maximum firing rate in a musculoskeletal model of human reaching

**DOI:** 10.1101/2025.09.18.677036

**Authors:** Tiina Murtola, Christopher Richards

## Abstract

Vertebrate muscle has two features which heavily influence its force generation characteristics: rate-coded control and the division of muscles into multiple motor units (MUs). Studying how these features affect functional outcomes is difficult in vivo, but also in silico due to scarcity of modelling frameworks integrating rate-coded MU pool models with musculoskeletal models for functional tasks. We implement a human upper limb reaching model with muscles consisting of multiple rate-coded MUs and demonstrate its ability to generate accurate reaching movements under online control. Using this model, we investigate how reaching performance is influenced by two key features of MU pools, the number of MUs and their maximum firing rates. Our simulations suggest that very small MU pools with low firing rates tend to produce less accurate reaching movements compared to larger pools with higher firing rates. Increasing either the size of the pool or the maximum firing rate generally improves performance, but benefits become negligible beyond about 10-20 MUs and firing rates of 25-50 Hz. This pattern holds for the entire workspace but targets on the distal boundaries appear more sensitive to MU pool properties. Hence, our results indicate that rate coding and MU properties may play a part in determining functional outcomes in a task- and MU pool-dependent manner.

## 1 Introduction

Generating and controlling movement relies on complex interplay between neural and muscular modulation of force generation. Understanding how the inherent characteristics of the neuromuscular system impact this modulation is central in motor control research and applications. Two salient features of force generation in vertebrate muscle are the rate-coded control (i.e. the modulation of force through the frequency of stimulation impulses [7, 31]) and the division of muscles into multiple motor units (MUs; [40]) of varied fibre types. Both features are known to vary across muscle types and species [32], but it is not known how the number of MUs and their firing rates influence functional outcomes.

Studying the fundamental relationship between the characteristics of multi-MU muscles and functional behaviour is challenging *in vivo*, although MU output features, such as the firing rates and impulse timings, have been studied in conjunction with simple behaviours such as low and medium force ramps [11, 19] or trapezoidal force profiles [3, 4, 10, 27], and explosive isometric contractions [12]. While such experimental studies provide valuable information about MU behaviour, tasks are often limited to isometric or near-isometric conditions particularly for techniques such as high-density surface electromyography [6], the MUs studied represent only a part of the entire muscle MU pool [13], and the number of confounding factors (including the control strategy employed by the central nervous system) is large. Furthermore, many underlying MU pool characteristics must be determined invasively [e.g. 2], making it difficult to deduce how these characteristics may affect both MU firing behaviour and functional outcome *in vivo*.

Computational MU pool models [e.g. 16, 35, 4] provide a tool for investigating how the distribution of MU characteristics within a muscle affect its output, but most studies focus on a single isolated muscle performing slow or constant, often isometric, contractions [e.g. 16, 20, 35, 36, 5], which provides limited insights into the whole organisms producing movements in a real environment. Musculoskeletal (MSK) models have the potential to provide this insight by providing a way to track causal chains from muscle excitations to contraction forces to whole-organism function [8, 39]. However, the muscle models typically used in MSK models lack the key features (multiple MUs and rate-coded control) needed to capture the dynamic behaviour of force generation in multi-MU muscles. Hence, our ability to address questions related to the functional role of these inherent muscle features is limited by lack of a framework enabling integration of the two modelling approaches.

The goal of the present work is to integrate rate-coded multi-MU muscles, which have been validated against high-density surface EMG data [34], into a MSK model of human reaching with predictive feedback control. We aim to investigate the fundamental principles governing movement generation with rate-coded multi-MU muscles. To achieve this, we study goal-oriented reaching, which is a behaviour with sub-maximal mechanical demands, to better understand how the number of MUs, *N*, and their maximal firing rate, *r*_*max*_, might limit performance across the workspace.

## 2 Methods

The simulations were carried out using a conceptual 3 d.o.f. horizonal reaching model [33] with six monoarticular muscles, each with properties corresponding to a young adult (Figure 1a). Two changes were made to the original model as detailed below: All muscles in the original model were replaced with muscles consisting of pools of MUs, and the predictive PD controller governing movement was updated to enable rate-control of the MU pools.

**Figure 1:**
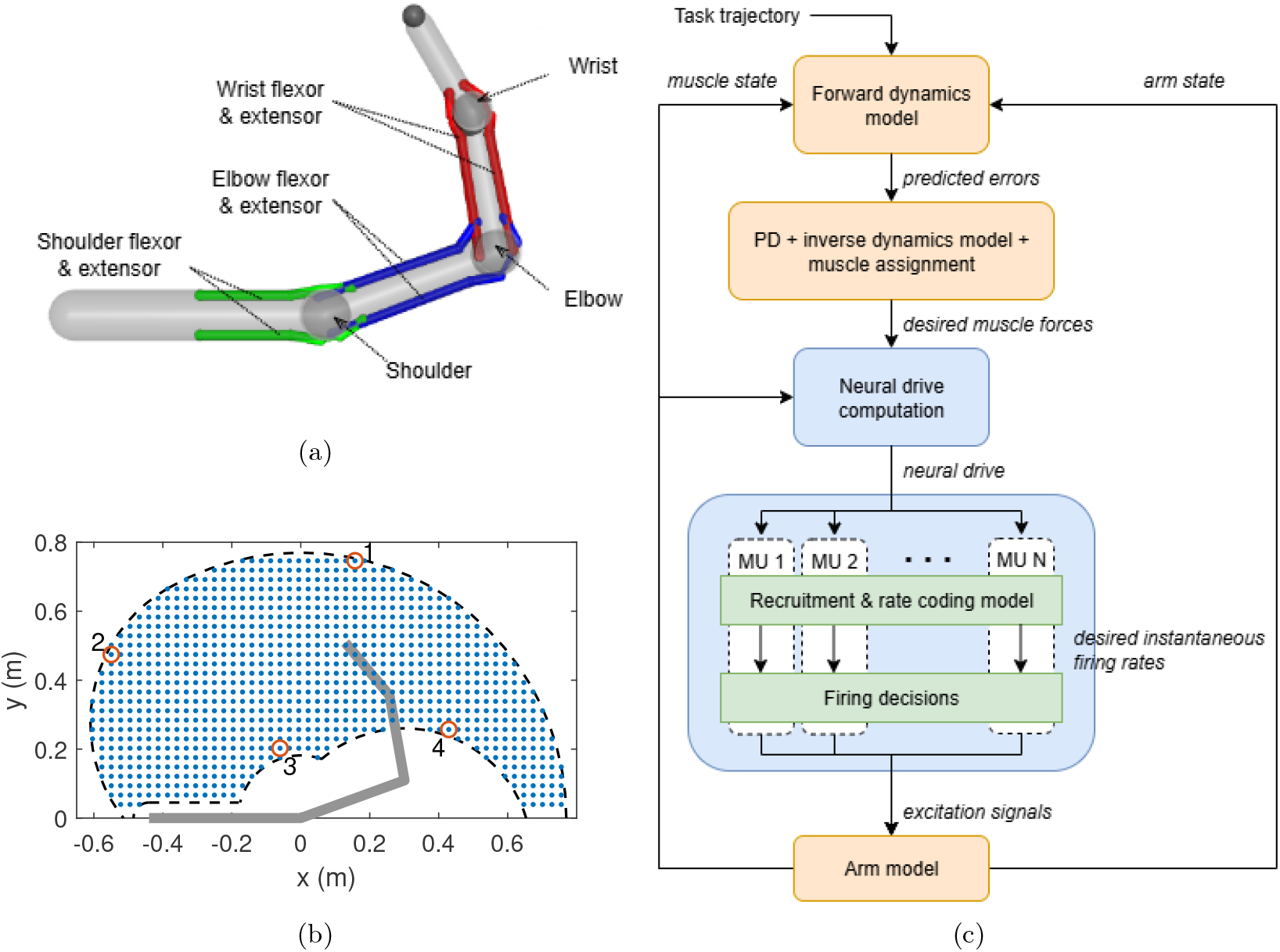
(a)) The three d.o.f. arm model with six monoarticular muscles. (b)) The optimisation targets numbered 1-4 (red circles) and a uniform grid of workspace targets (blue dots) relative to the boundaries of the workspace (dashed black) and initial arm position (thick gray line). (c) The control scheme for the reaching model, with the original control structure [33] in yellow and the new blocks for neural drive computation and recruitment and rate coding shown in blue. Note that the muscles in both the arm model and the forward dynamics model consist of multiple MUs in the present work but this does not affect the state information flow.

The MU pool model used was based on a model with mixed-linear-exponential recruitment threshold, a logarithmic rate function [34] which approximates the behaviour of a leaky-integrate-and-fire model [3], and a uniform *r*_*max*_ for all MUs in the pool. This MU pool type was observed to perform slightly better than alternative recruitment and rate coding models when matching reaching forces, with constant *r*_*max*_ for all MUs performing more consistently across different targets than onion-skin type *r*_*max*_ schemes [34]. In order to use the same general MU pool structure for the six muscles despite their different isometric strengths, the MU pool equations were rewritten using normalised parameters. This also enables comparison among pools with different numbers of MUs and maximum firing rates. The model equations can be found in the Appendix.

The number of MUs in real upper limb muscles is too large to feasibly allow the necessary optimisations and repeated simulations for the present study. Instead, we use smaller MU pools constructed to represent baseline property distributions in large (*N* ≥300) pools (Figure 2). This was done by calculating equivalent distribution parameters which preserved the shapes of the distribution functions regardless of pool size *N*. The details of the calculations and the parameter values are given in the Appendix.

**Figure 2:**
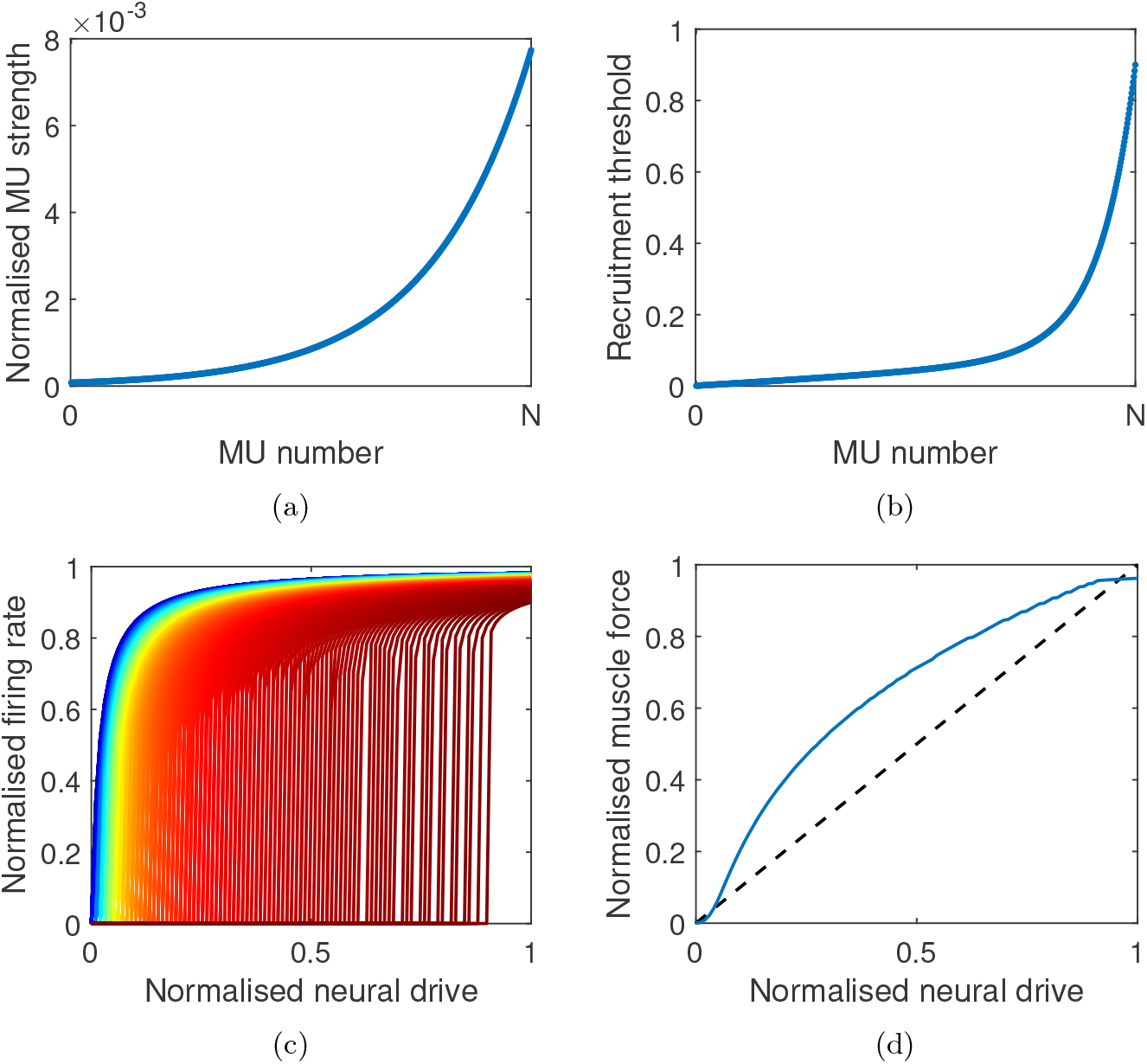
The properties of the large baseline MU pool representing upper limb muscles. (a) Distribution of MU strengths normalised with total muscle strength against MU index from 1 to *N*. (b) Recruitment threshold (in terms of normalised neural drive) vs MU index. (c) Firing rates normalised with *r*_*max*_ vs neural drive normalised with total muscle strength. MU index is indicated by colour (first MU = blue, last MU= dark red; only every 10^th^ MU shown). (d) Steady state tetanic muscle force vs neural drive, both normalised with total muscle strength. One-to-one correspondence (dashed) is shown for reference.

The control scheme of the reaching model [33] was used to control movements but with the inverse muscle model replaced with a recruitment and rate coding model. To summarise this scheme (Figure 1c), the controller utilises a forward dynamics model to predict position and velocity errors with respect to a pre-planned hand trajectory. These errors are used to calculate desired joint torques, which are converted to desired muscle forces. These signals are then passed to the recruitment and rate coding model, which combines them with predicted force-velocity-length state of the muscle to calculate the normalised neural drive signal for the MU pool. This signal drives threshold-based recruitment of the MUs, and, once recruited, determines the instantaneous firing rate of each MU. Finally, the desired inter-impulse-time (inverse of firing rate) for each MU is compared with time elapsed since its last excitation impulse to determine whether an impulse should be triggered. If an impulse is triggered, the excitation signal for the corresponding MU is set to 1 for 3 ms (otherwise excitation signals are zero), and the excitation signals are then used as inputs to the muscles of the arm model, which converts them to an activation level through a dynamic third order model [28]. Resulting changes in muscle contraction state and arm configuration are passed to the controller’s forward dynamics model, allowing it to update error predictions for the next time step. When calculating predicted errors, the forward model assumes no further control actions are taken, but ongoing excitation impulses will be completed.

Simulations were carried out with 49 different MU pool model variants. These variants were the combinations of seven pool sizes, *N* = 1, 2, 5, 10, 20, 50, and 100, and seven maximum firing rates, *r*_*max*_ = 20, 25, 30, 40, 50, 75, and 100 Hz. For each model, control parameters were optimised by minimising the average homing-in error, *e*_*hi*_, over four targets. The homing-in error calculates the average error in position from the moment when the planned trajectory reaches the target to the end of simulation, and it captures the accuracy of the reaching movement. The four *optimisation targets* are located at the boundaries of the arm model’s workspace, and they have previously been used to obtain control parameter sets usable for the entire workspace [33]. A second set of simulations covering the entire workspace (using a *full workspace target set*) are also performed using the model-specific optimised control parameters (Figure 1b).

The arm model is implemented in MuJoCo (2.0), with custom code for the muscles and control models. The control parameter optimisation is carried out with Matlab (R2023a Update 3, with Global Optimisation Toolbox version 4.8.1) using genetic algorithm with default settings, and optimisations are repeated to minimise the chance of local optima. All models, parameter sets, and codes are available from [repository link tbc].

## 3 Results

Performance at reaching for the optimisation targets depends on both the number of MUs, *N*, and the maximum firing rate, *r*_*max*_, but also on the target location. This is exemplified in Figure 3 with four example models: 1 and 20 MUs with *r*_*max*_ of 20 and 50 Hz. Overall, the homing-in error is high when both *N* and *r*_*max*_ are low, and decreases when either parameter is increased (Figure 3a). However, the relative magnitudes of the effects are target-specific. For example, performance for target 2 (contralateral) is improved notably more by increasing *r*_*max*_ from 20 Hz to 50 Hz compared to increasing *N* from 1 to 20, while for the other targets increases in either firing rate or motor unit number leads to similar improvements.

**Figure 3:**
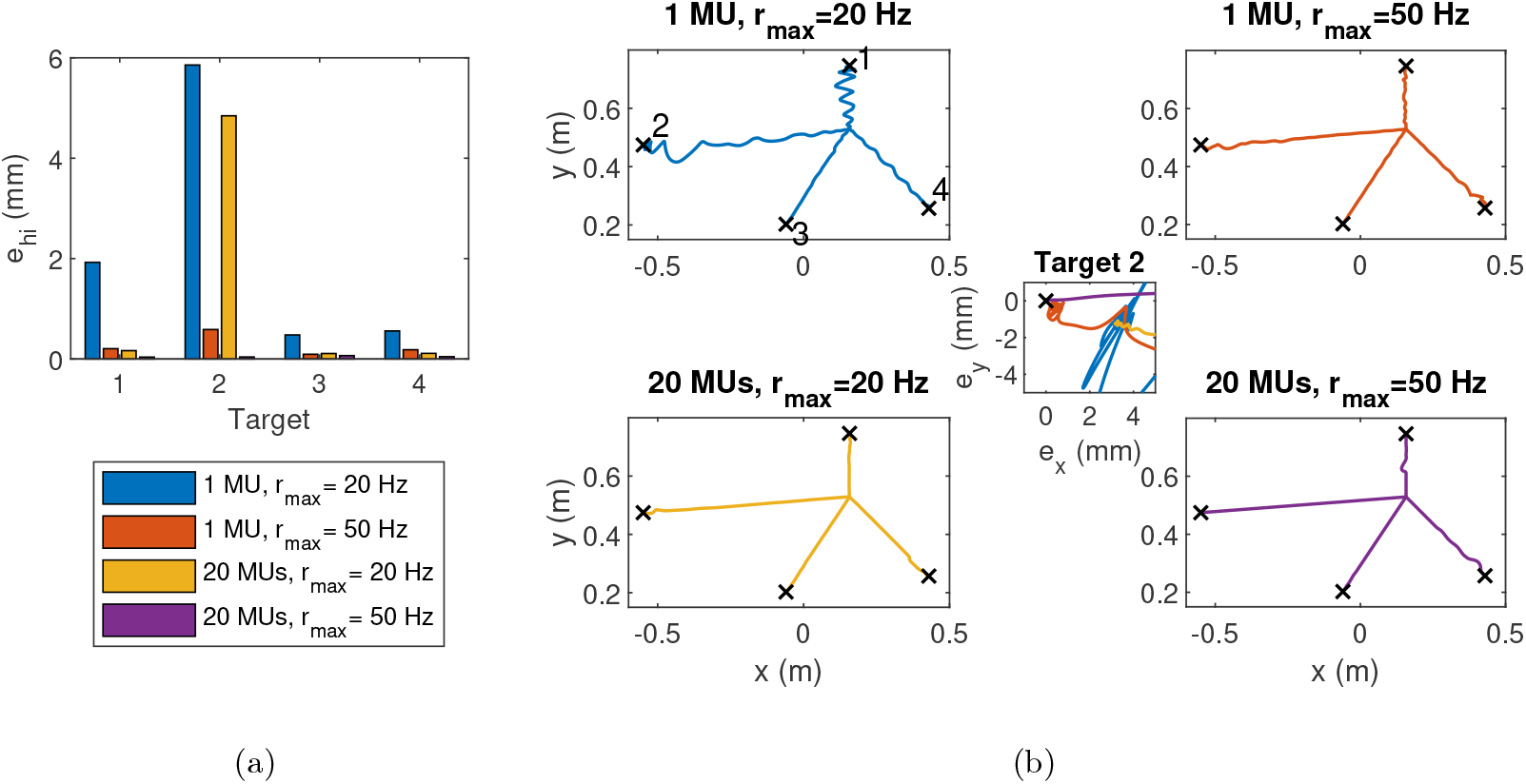
The trajectories and homing-in errors for four example models. (a) Homing-in errors (*e*_*hi*_) for the four models grouped by target. (b) Each of the main panels shows the trajectories to the four targets (black crosses) for one of the four models: 1 MU and 20 MUs each with *r*_*max*_ values of 20 and 50 Hz. The first panel also shows target index. The middle insert shows a zoomed-in view of target 2 with the final stages of trajectories for the four models. Position in the insert is shown as error relative to the target location.

In addition to the directly optimised homing-in error, the MU pool also affects the quality of the movement, including the the straightness of the trajectory to target. A low number of MUs combined with low maximum firing rate results in visibly non-straight trajectories, especially for the forward and contra-lateral targets (Figure 3b). The straightness of the trajectory increases with both number of motor units and with maximum firing rate, although individual “bumps” may still occur, for example, when a joint reaches the end of its range of motion. After 20 MUs with 50 Hz *r*_*max*_ the general improvement in trajectory straightness is no longer discernible.

Although performance at individual targets can vary, the overall trend of performance increasing with the number of MUs or the *r*_*max*_ holds when considering the average homing-in error across the four optimisation targets (Figure 4). This performance gain is most notable when comparing very small pools of MUs (*N* ≤10) and when the maximum firing rate is is low to medium (*r*_*max*_ ≤50 Hz). Once either *N* or *r*_*max*_ gets sufficiently high, the performance gain obtained by increasing the other variable becomes negligible. One clear exception to this is that the average homing-in error decreases notably when *r*_*max*_ is increased from 20 to 25 Hz regardless of the pool size. This phenomenon is largely driven by performance at target 2 (contralateral), where at very low firing rates the large oscillations along the trajectory result in the arm not quite reaching the target (Figure 3A).

**Figure 4:**
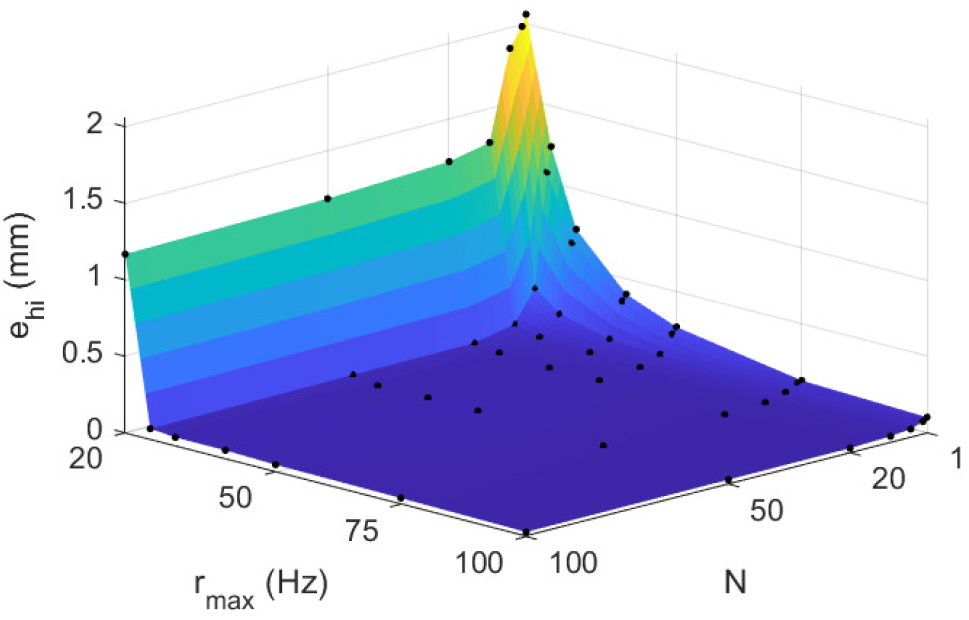
The average homing-in error for the four optimisation targets for different numbers of MUs and *r*_*max*_ values. Black dots indicate values obtained from simulations, and the surface between them is obtained through linear interpolation.

Simulations of targets across the entire workspace reveal that patterns observed for the optimisation targets are broadly valid for most targets, even when performance has not been optimised for them. For most targets in the workspace, the homing-in error surfaces against *N* and *r*_*max*_ (Figure 5, small panels) are similar to Figure 4, with the lowest errors at larger pools with higher *r*_*max*_. Target location mainly affects error magnitude for the worst performing models, and whether increasing *N* or *r*_*max*_ is more effective in improving performance relative to the *N* = 1, *r*_*max*_ = 20 Hz model. Note, however, that a few targets at the distal workspace boundaries exhibit more complex behaviour with local minima and maxima in the error surface. To summarize the difference and variation in *e*_*hi*_ between the models, the main panel Figure 5 shows the (log-scaled) standard deviation calculated across all models. The results suggest that while larger pools with higher *r*_*max*_ are beneficial across the entire workspace, largest performance losses and gains were seen near the distal workspace boundaries when the MU pool composition was varied.

**Figure 5:**
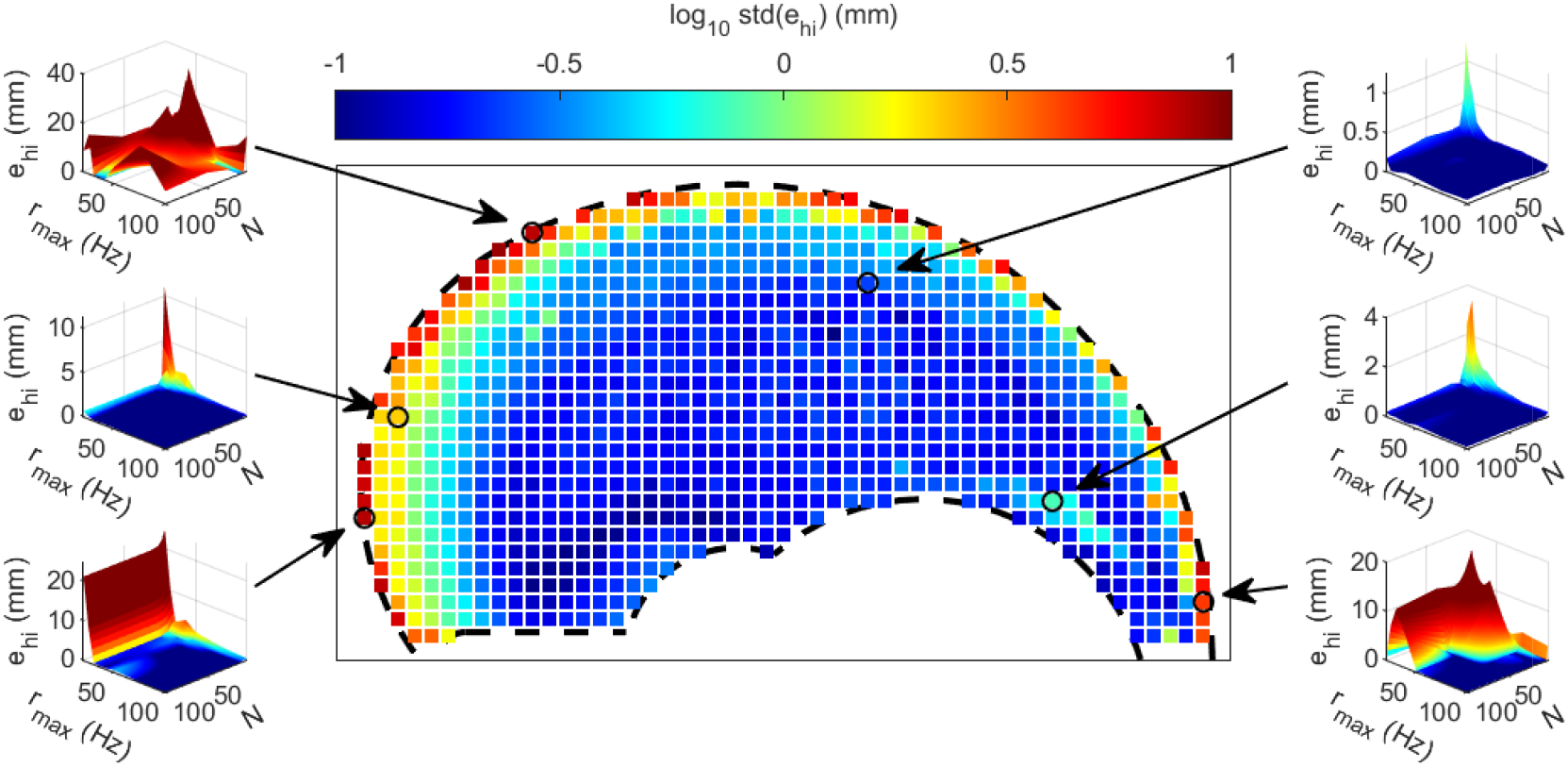
The variability (standard deviation) of the homing-in error across all models over the entire workspace. Each dot indicates a target and its colour the standard deviation value (log-scale), with the workspace boundaries indicated by dashed lines. Small panels show the source of this variability by displaying the homing-in error vs *N* and *r*_*max*_ for selected targets (indicated by arrows and circles). The scale of the y-axis in the small panels is linear, but surface colours match the log-scale of the middle panel and colourbar.

The median homing-in error value across the entire workspace is displayed in Figure 6, with the two panels showing the same data using different groupings. In models with the same number of MUs, increasing *r*_*max*_ leads to lower performance error until *N* = 10 (Figure 6 top panel). If *N* ≥ 20 and *r*_*max*_ ≥ 25 Hz, further increases in parameter values do not necessarily produce improvements. The picture is slightly more mixed when considering the impact of *N* on pools with constant *r*_*max*_ (Figure 6 bottom panel). Although increasing the number of MUs is generally beneficial, at some *r*_*max*_ values larger pools can result in higher errors than a slightly smaller pool (e.g. 2 MUs performs worse at 20 and 25 Hz than 1 MU, and 50 MUs worse than 20 MUs at 40 Hz). Overall, Figure 6 indicates that the pattern observed in Figure 4 for the optimisation targets holds generally across the workspace.

**Figure 6:**
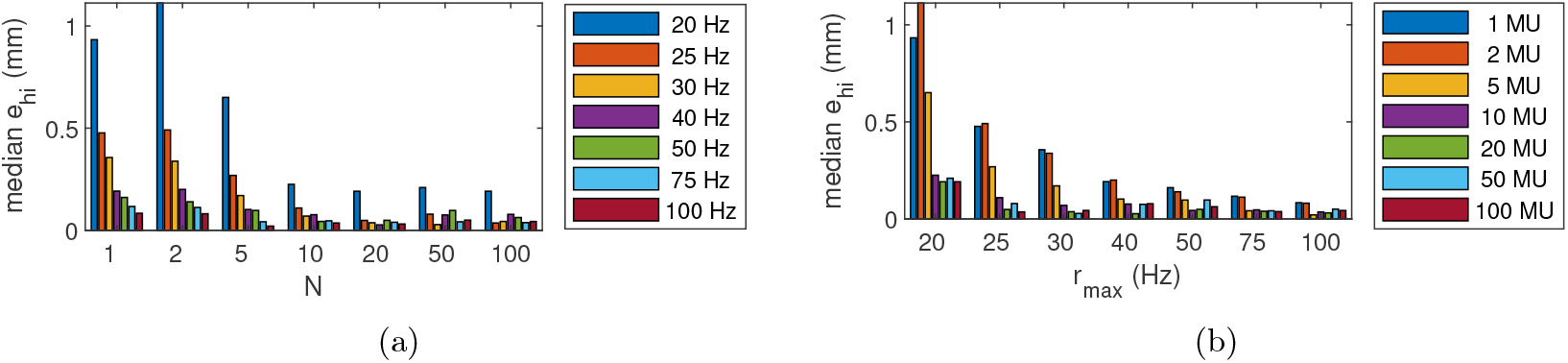
Median homing-in error (*e*_*hi*_) across the entire workspace. (a) Models have been grouped by number of MUs (*N*) with colours indicating maximum firing rate (*r*_*max*_). (b) The same data but grouped by *r*_*max*_ with colours indicating *N* .

## 4 Discussion

In the present work, we have integrated rate-coded multi-MU muscles into a MSK model of human reaching in order to investigate how the fundamental features of MU pools affect movement generation. Our results suggest that rate-coded multi-MU muscles can be used in MSK models to successfully generate reaching movements using online control, but the number of MUs and their maximum firing rates can have an impact on the accuracy and trajectory of reaching. In particular, very small MU pools with low firing rates tend to struggle to follow the planned straight trajectory to the target and end up with higher final position errors compared to larger pools with higher firing rates. This pattern holds generally for targets across the entire workspace, but targets near the distal boundaries of the horizontal workspace are particularly sensitive to changes in MU pool properties.

The relatively poor performance of small MU pools with low maximum firing rates stems mainly from the fact that rate-coding quantises motor control; force can only be produced in “units” of twitches. When the number of MUs is small, this quantised control can lead to reduced performance due to two reasons. First, because the total muscle strength is the same for all MU pool models, MUs in small pools have higher isometric strengths and twitch amplitudes, making small corrective movements more likely to result in overshooting. While this effect could have been mitigated lowering twitch amplitudes, changes in twitch properties impact multiple aspects of force production (e.g. total muscle strength and the speed at which force can be ramped up) simultaneously, potentially resulting in other problems. Second, multi-MUs with different recruitment thresholds increases the likelihood of asynchronous initiation of twitches within a muscle, smoothing out total muscle force which results in smoother joint movements which are less likely to trigger series of overcorrections. This effect is particularly notable with the smallest MU pools, where adding just few extra MUs improves the ability of the MU pool to generate the smooth force profile, similarly to experimentally observed smoothing of contractions via asynchronous stimulation in cat soleus muscles [37]. Notably, in our simulations this happens even though the controller does not intentionally implement asynchronisation. Low maximum firing rate impacts performance through similar routes as the number of MUs; reducing *r*_*max*_ increases twitch amplitudes (to preserve tetanic force) and it also limits the speed at which force production can be ramped up.

Our goal was to obtain fundamental understanding of the relationship between MU pool features and function by constructing theoretical MU pools. However, to provide the simulation results context, let us compare the pools to known features of human upper limb muscles. Real human muscles tend to have a few hundred MUs [14, 1]. Hence, even though the number of MUs can decrease, for example, due to ageing [42] or neurodegenerative diseases [24, 17, 29], or after stroke [30, 23] or spinal injury [41], only a very dramatic loss of MUs would breach the limit (*N* ≈10–20) where the number of MUs starts to notably impact reaching performance for most targets. Hence, loss of MUs due to the above pathologies would be unlikely to impact a task such as reaching (though the impact on more demanding tasks requires further study).

In contrast, changes in firing rate may be more consequential during real behaviour. Maximum firing rates reported in human upper limb muscles during slowly varying or sustained force tasks have been reported to range from 18 [38] to 29 Hz [11], with a higher level of contraction associated with higher maximum firing frequency [11]. These values are on the low end of the *r*_*max*_ values used in simulations, although human MUs can exhibit higher firing rates, for example, during rapid contractions [12, 15], to temporarily enhance force produced during submaximal contraction or to mitigate force loss due to fatigue [15]. In the model, *r*_*max*_ is a hard upper limit, and in most reaching simulations it is the peak firing rate reached by the first-recruited MUs. Hence, the relevant comparison is between observed firingn rates during similar task conditions in simulations versus experiments. In these cases, the MU pool models with low maximum firing rates (*r*_*max*_ ≤ 30 Hz) are most relevant for slow, low-force movements such as reaching. At these firing rate levels, models exhibit sensitivity to both *r*_*max*_ and *N* suggesting that a larger MU pool may be needed to compensate for the limitations placed by the low firing rates at which MUs often operate.

While our simulations covered the entire horizontal workspace of the model, this represents only a fraction of the range of upper limb movements humans utilise. Hence, even though our results suggest that only a very small number of MUs is necessary for the tasks studied, the significantly higher number of MUs in real muscles could convey benefits in other types of tasks. Furthermore, motor performance is inherently dependent on the control strategy employed. In our model, key elements of this control strategy are modelled after known features of real MU pools: a common neural drive [9] and threshold-based recruitment that follows the size principle [18]. While the real neuromuscular system can utilise a more nuanced control approach [21], our model cannot deviate from these principles, which can lead to badly timed or suboptimally ordered recruitment of MUs relative to task demands. The current control strategy is likely sufficient for the slow, low-force reaching movements in the present study, but future studies on how MU pool characteristic impact performance in a larger range of tasks will likely need to also address the generalisability of the control strategy, for example, utilising synergies [21] or recruitment based on factors such as metabolic cost [26].

Our results suggest that the distal boundary of the workspace is particularly sensitive to the properties of the MU pools. This effect may be partially attributable to the naive controller, which does not include an effective strategy for dealing with singularities in the arm position or joints reaching their maximum range of motion. However, despite these challenges related to arm position, the controller seems generally more able to navigate these challenging reaches when the MU pool is larger or *r*_*max*_ is higher. While some of these challenges are specific to the model, the reaching targets at the distal workspace boundary is also challenging in reality, requiring precise control of an extended arm using proximal muscles. Hence, the possibility that larger, high firing frequency pools require less sophisticated motor control strategies warrants further investigation.

While our simulations link MU pool properties and accuracy of reaching, it is worth noting that the majority of the workspace could be reached with final position errors below 1 mm regardless of which MU pool was used. From the perspective of daily movements, this accuracy is often sufficient; hence differences in performance may have limited practical significance. The different MU pools likely would, however, perform differently in movements where accuracy is not the sole performance metric of interest. For example, while not presented in the current paper, small MU pools with low maximum firing rates tend to have higher co-contraction levels during reaching because large twitch amplitudes and nonlinear summation of twitches can make smaller joint torques easier to generate at high force levels. Since co-contraction can increase inefficiency in a noise-free system but be an energy-efficient response to expected noise or uncertainty [22], the net impact of differences in MU pool size or firing characteristics likely depends heavily on the performance factors considered and how they are weighed in the optimisation cost function. A systematic investigation into how different MU pool characteristics impact different aspects of performance is, however, left for future studies.

Our simulations indicate that rate-coded MU-pools can be integrated into MSK simulation frameworks to simulate successful reaching movements, but that the characteristics of the MU pools may impact performance outcomes. The performance loss observed in the case of small MU pools and low *r*_*max*_ values could be considered to arise mainly from a mismatch between force profiles required by the task and the profiles the MU pool can feasibly produce, similarly to the challenges experienced by MU pools in force matching outside MSK simulations [34]. This, combined with our observation of target-dependent variation in performance, suggests that best functional outcomes might be reached when the force production characteristics of the MU pool matches the requirements of the tasks. In this light, our results suggest that force production in larger MU pools with higher maximum firing rates confer more versatile force production to enable a larger range of tasks.

## Acknowledgments

This research was funded by the Wellcome Trust Investigator Award 215618/Z/19/Z.

## Appendix

### Multi-MU muscle model

As in the original model [34], the total force generated by muscle *j* (*j* = 1, …, 6), *F*_*j*_(*t*), at time *t* produced by a pool of *N* MUs is

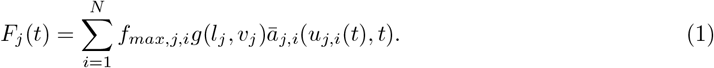

The force produced by the *i*^th^ MU (*i* = 1, …, *N*) is a product of its isometric strength, *f*_*max,j,i*_, FVL gain *g*(*l*_*j*_, *v*_*j*_) depending on the length *l*_*j*_ = *l*_*j*_(*t*) and contraction speed *v*_*j*_ = *l*·_*j*_ of the muscle, and the normalised activation state ā_*j,i*_(*u*_*j,i*_(*t*), *t*) of the MU arising from the excitation signal *u*_*j,i*_(*t*). The MUs are ordered from smallest (*i* = 1) to largest (*i* = *N*), with the total isometric muscle strength *F*_*max,j*_ distributed exponentially among the MUs [34, 16]:

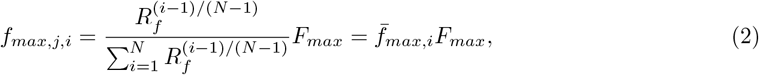

where *R*_*f*_ = *f*_*max,N*_ */f*_*max*,1_ is the range of MU strengths, *F*_*max*_ is the isometric strength of the entire muscle and 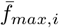 is the normalised MU strength. The speed of the activation dynamics ā_*j,i*_, which is governed by three time constants, is assumed to scale with the strength of the MU following [16]. The time constants of the third order activation model [28] are scaled with

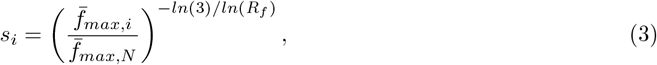

based on [16], which results in equivalent scaling of the time to peak and half relaxation time of twitches. The baseline time constants (i.e. for the *N* ^th^ MU) are *α* = [18.5, 31, 14] ms and the non-linearity constants *β* = [0.6, 0.8, 0.7], resulting in time-to-peak of 57.5 ms and half-relaxation time of 64 ms for the fastest MU.

The excitation signal *u*_*j,i*_(*t*) is the outcome of recruitment and rate coding. The *i*^th^ MU of the *j*^th^ muscle is recruited when a normalised (dimensionless) neural drive Ī_*j*_ (*t*) = *I*_*j*_(*t*)*/F*_*max,j*_,where *I*_*j*_(*t*) is the raw neural drive from the PD controller assigned to muscle *j*, exceeds the relative (dimensionless) recruitment threshold 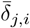 defined as

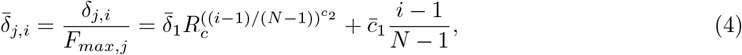

where 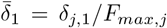 and 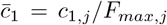, and *δ*_*j,i*_ is the threshold for the original model [34, 3] defined using unnormalised parameters *δ*_*j*,1_, *c*_1,*j*_ and *c*_2_. Hence, we assume that relative to the strength of the muscle, the thresholds of all MUs in the six muscles of the arm model are identical.

Once the *i*th MU of the *j*th muscle has been recruited, its firing rate follows the rate function

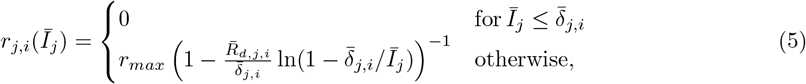

where *r* _*max*_ is the maximum firing rate (assumed equal for all MUs and muscles), 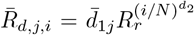 with normalised parameter 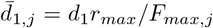, and *d*_1_, *d*_2_, and *R*_*r*_ are parameters as in the original model [34]. ln() denotes the natural logarithm. For the normalised model, the 90% rate coding band limit (relative to *F*_*max,j*_), that is value of Ī_*j*_ at which *r*_*j,i*_ = 0.9*r*_*max*_, is

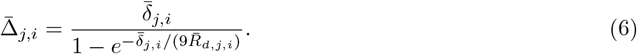

The strategy for constructing *u*_*j,i*_(*t*) from *r*_*j,i*_(*t*) [34] is implemented as a real-time algorithm to enable the reaching controller to decide whether to trigger an excitation impulse for any MUs at every time step, based on the comparison of 1*/r*_*j,i*_(*t*) and time elapsed since last excitation impulse. The only modification made to the original algorithm is that the impulse duration was made independent of simulation timestep and fixed to 3 ms.

### Constructing baseline and equivalent MU pools

In order to construct small MU pools that can be used to represent upper limb muscles, we first define a “full” MU pool. The shapes of the MU pool characteristics and behaviours (Figure 2) are not sensitive to the exact number of MUs when the number is large (*N* ≥ 300), which is typically the case for human limb muscles (e.g. *N* ≈300 for biceps brachii [14]). Since the muscles in the reaching model represent entire muscle groups, we select *N*_*full*_ = 600, which also ensures that the steady-state response *F*_*j*_(*I*) is smooth. A wide recruitment range was used with 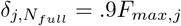, as this is typical for human upper limb muscles [11, 25]. The first recruitment threshold, *δ*_*j*,1_ = 0.001*F*_*max,j*_, was set by trial and error to be well above the amplitude of a single twitch but low enough to ensure recruitment of MUs even in low force tasks. The other parameters values for the baseline pool are given in Table 1.

**Table 1:**
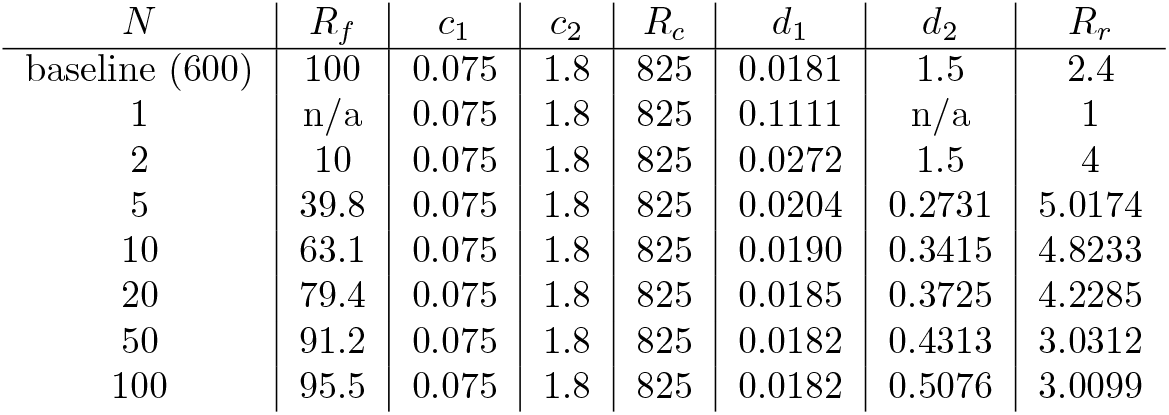
MU pool parameters for the baseline (not used for computations) and the seven pool sizes investigated. Baseline parameters are based on [34], except *R*_*c*_, which is calculated from threshold parameters.

To construct MU pools that preserve the shapes of the MU property distributions when *N << N*_*full*_, we note that the distribution functions can be re-written using a normalised MU index

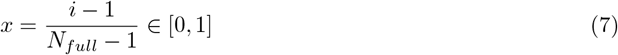

which can be considered approximately continuous given the large *N*_*full*_. Note that the large *N*_*full*_ also allows writing the rate function Eq. 5 in terms of *x* without adjustment of parameter values.

For a small pool with *N* MUs, the MU strengths are calculated by partitioning *x* into *N* equal sections with the boundaries between sections denoted *x*_0_, *x*_1_, …, *X*_*N*_. The average *f*_*max*_(*x*) each section is obtained via integration from *x*_*i*−1_ to *x*_*i*_, and the values obtained for the first and last segment are used to calculate an equivalent *R*_*f*_ for the small pool,

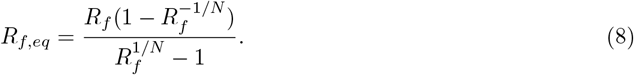

The *f*_*max,i*_ values are then be calculated from Eq. 2 using *R*_*f,eq*_ instead of *R*_*f*_. This will automatically ensure that MU strengths add up to *F*_*max*_ and that as *N* → *N*_*full*_, the small pool distribution converges to the full pool distribution.

Recruitment thresholds and the rate function are more challenging, as in addition to preserving the shape of the functions, the resulting *F* (*I*) should also match the full pool steady state response. To achieve this, the partitioning of *x* into *N* segments is again used. The recruitment threshold value at *x*_*i*−1_ (*i* = 1, …, *N*) is assigned as the recruitment threshold for the *i*^th^ MU in the small pool. The 90% rate band limit at *x*_*i*_ is assigned as the equivalent limit for the *i*^th^ MU in the small pool, and its value is used to calculate the rate function parameters from Eq. 6 by utilising the first, last and and middle MUs. Specifically, when *N* = 1, 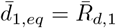while other parameters are irrelevant; when *N* = 2, *d*_1,*eq*_ as before, 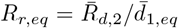, and *d*_2_ is kept at default; and when *N* ≥2, *d*_1,*eq*_ and *R*_*r,eq*_ as before and *d*_2_ is calculated from the middle MU. Alternative ways of calculating the rate parameters, including using the threshold and rate band values to calculate MU-specific parameters, were explored but their impact on overall MU pool characteristics was not large enough to justify the added complexity. The resulting parameter values are listed in Table 1 for each MU pool size investigated.

## Notes

### Competing Interest Statement

The authors have declared no competing interest.

